# Landscape genomics of an obligate mutualism: discordant population structures between a leafcutter-ant and its fungal cultivars

**DOI:** 10.1101/458950

**Authors:** Chad C. Smith, Jesse N. Weber, Alexander S. Mikheyev, Flavio Roces, Martin Bollazzi, Katrin Kellner, Jon N. Seal, Ulrich G. Mueller

## Abstract

To explore landscape genomics at the range limit of an obligate mutualism, we used genotyping-by-sequencing (ddRADseq) to quantify population structure and the effect of host-symbiont interactions between the northernmost fungus-farming leafcutter ant *Atta texana* and its two main types of cultivated fungus. At local scales, genome-wide differentiation between ants associated with either of the two fungal types is greater than the differentiation associated with the abiotic factors temperature and precipitation, suggesting that specific ant-fungus genome-genome combinations may have been favored by selection. For the ant hosts, we found a broad cline of genetic structure across the range, and a reduction of genetic diversity along the axis of range expansion towards the range margin. In contrast, genetic structure was patchy in the cultivated fungi, with no consistent reduction of fungal genetic diversity at the range margins. This discordance in population-genetic structure between ant hosts and fungal symbionts is surprising because the ant farmers co-disperse with their vertically-transmitted fungal symbionts, but apparently the fungi disperse occasionally also through between-nest horizontal transfer or other unknown dispersal mechanisms. The discordance in population-genetic structure indicates that genetic drift and gene flow differ in magnitude between each partner in this leafcutter mutualism. Together, these findings imply that variation in the strength of drift and gene flow experienced by each mutualistic partner affects adaptation to environmental stress at the range margin, and genome-genome interactions between host and symbiont influences adaptive genetic differentiation of the host during range evolution in this obligate mutualism.

## 1 Introduction

Range expansions can be impeded or facilitated by mutualistic or competitive interactions between species (Gilman et al. 2010; Lavergne et al. 2010; Urban 2011). In mutualisms, codependencies between mutualistic partners can facilitate range expansion, for example when association with a symbiont increases the niche breadth under which a host can exist (Afkhami et al. 2014; Douglas 2009; Nobre et al. 2010). Mutualisms can also impede range expansion, for example when a symbiont tolerates a narrower window of environmental conditions (e.g., temperature) compared to the conditions tolerated by a host (Bronstein 1989; Dixon et al. 2015; Hume et al. 2016), and thus the range of a host is determined by its symbiont. Generally, the importance of species interactions for either facilitating or restricting range adaptations of a host are less understood than range-limiting adaptations driven by abiotic range-limiting factors, such as temperature or precipitation (Sexton et al. 2009; Urban 2011; Schoville et al. 2012).

Much work on the evolution of range-limiting species interactions has been theoretical (Case et al. 2005; Lavergne et al. 2010; Gilman et al. 2010; Holt et al. 2011; Urban 2011; Norberg et al. 2012), primarily because it is difficult to test adaptation to range-limiting factors (Holt 2003; Hoffmann & Sgrò 2011; Angert et al. 2011). Empirical work has focused mostly on range-limiting competitive and antagonistic interactions (Cunningham et al. 2009; Hellmann et al. 2012; le Roux et al. 2012), and less so on range-limiting mutualisms, such as plant-pollinator, plant-microbe, or insect-endobacteria mutualisms (Bronstein 1989; Bronstein & Patel, 1992; Smith et al. 2011; Stanton-Geddes & Anderson 2011; Thompson & Rich 2011; Moeller et al. 2012; Moran 2016). Adaptations that alter range limits are least understood for mutualisms with vertical transmission of a symbiont (e.g., plant-endophyte or insect-endobacteria mutualisms), where the two partners function as an integrated unit, and fitness of the association is determined by genome-by-genome interactions (inter-genomic epistasis; Wolf et al. 2000; Heath 2010) and sometimes by complex host-symbiont interactions, such as host-symbiont conflict (Mueller et al. 2005; Wade 2007).

As populations expand into new habitat, the action of selection, drift, and gene flow can influence adaptive potential and leave a distinct footprint on the genome (Sexton et al. 2009, 2014; Excoffier et al. 2009). Successive founding events at an expanding range edge, for example, can result in strong genetic drift over a large geographic area, reducing both genetic diversity and efficacy of selection as new populations at the range edge adapt to novel environmental conditions. Gene flow from the interior of the range can reconstitute genetic diversity lost in populations at the range edge, but gene flow can also swamp out adaptive variation accumulating at range limits (Kirkpatrick & Barton 1997; Holt 2003). In a mutualism, the evolutionary forces acting during a range expansion could be similar or different for each mutualistic partner, potentially affecting the colonization of new habitat.

Here, we examine a range expansion of an ant-fungal mutualism to evaluate the effect of host-symbiont interactions on genetic differentiation relative to the effects of two abiotic factors, temperature and precipitation (Figure S1). We couple genotyping-by-sequencing (ddRADseq; Peterson et al. 2012) of the leafcutter ant *Atta texana* with microsatellite-marker genotyping of *Leucocoprinus gongylophorus* fungi cultivated by the ants for food, to elucidate how range expansion affects population structure in each mutualistic partner and genome-wide differentiation in the ant host. We expected population-genetic structure to be similar between the ants and the fungus if the dominant evolutionary processes operating during the range expansion are similar for each partner in the mutualism, as might be predicted for the obligate leafcutter mutualism where the life cycles of the mutualistic species are inextricably linked and the ants co-disperse with their fungal cultivars. We also expected ant-fungus genotype combinations and climate to be correlated with genome-wide differentiation if symbiotic interactions and abiotic factors influenced evolution during the range expansion.

## 2 Materials And Methods

### 2.1 Study system

The leafcutter ant *Atta texana* is the northernmost representative of its genus (Bacci et al. 2009), ranging from the US-Mexico border region in north-east Mexico to northern Texas and western Louisiana west of the Mississippi River (Figure S1; Mueller et al. 2011a,b). The closest relatives of *A. texana* are the Mexican *Atta mexicana* and the Cuban *Atta insularis* (Bacci et al. 2009). The three *Atta* species diverged from each other before the last glaciation, presumably in southern North America. Following the end of the last glaciation about 11000 years ago, *A. texana* is thought to have expanded northward from ancestral populations in north-eastern Mexico (Mueller et al. 2011a,b). The first published observations of *A. texana* in Louisiana and eastern Texas documented a widespread presence of *A. texana* in counties at or near the current north-eastern range limits in Louisiana (Jones 1917; Snyder 1937; Walter et al. 1938; Smith 1939). Moreover, the current range limits of *A. texana* in Louisiana (Dash 2004; Mueller et al. 2011a,b) are essentially the same limits that were already recognized 80 years ago by Smith (1939), indicating that range expansion of *A. texana* was halted towards the east by the shallow water table of the Mississippi River basin (nests drown in areas of regular flooding) and halted towards the north by cold winter temperatures (Mueller et al. 2011a). This combined historical-biogeographic information suggests a conservative estimate for the north-eastward expansion of *A. texana* from southern populations sometime between 11000 and 100 years ago.

*A. texana* leafcutter ants cultivate a fungus called *Leucocoprinus gongylophorus* (Mueller et al. 2017; Mueller et al. 2018) which is obligately dependent on the ants and clonally propagated within nests and from maternal to offspring nests (Mikheyev et al. 2006, 2010; Mueller et al. 2010; Marti et al. 2015). *L. gongylophorus* is polyploid (multiple genomes per nucleus) and multinucleate (multiple nuclei per cell), with an observed average of 9.4 nuclei per cell for one fungus from *A. texana* (range 3-21 nuclei/cell; Carlson et al. 2017). The exact ploidy of *L. gongylophorus* is not known (most likely 3-7; Kooij et al. 2015; Carlson et al. 2017), ploidy appears variable between *L. gongylophorus* strains, and the number of nuclei per cell is variable in a mycelium (range 3-21 nuclei/cell; Carlson et al. 2017). Although *L. gongylophorus* clones are occasionally transferred between nests of sympatric leafcutter ant species through little-understood mechanisms (Mueller 2002; Mikheyev et al. 2006, 2007, 2010; Mueller et al. 2017), each leafcutter nest appears to cultivate only a single *L. gongylophorus* clone (Mueller et al. 2010). Population-genetic analyses of *L. gongylophorus* from *A. texana* found two main genotype clusters of cultivated fungi (so-called M- and T-fungi), which are distributed in sympatry across the range of *A. texana* (Mueller et al. 2011b). Because genetic admixture between M- and T-fungi occurs at low frequency (Mueller et al. 2011b), these two genotype clusters represent distinct clone-lineages of the same fungal species *L. gongylophorus*.

*A. texana* does not exist sympatrically with other leafcutter species throughout its range in Texas and Louisiana, so *L. gongylophorus* cultivars cannot be exchanged between different host species, whereas at other locations in the leafcutter range different sympatric leafcutter ant species can exchange cultivars between nests (Silva-Pinhati et al. 2004; Mikheyev et al. 2007; Mueller et al. 2017; Mueller et al. 2018). Absence of sympatric leafcutter species therefore simplifies analyses of ant-fungus co-evolution in *A. texana*, including analysis of possible species-specific adaptations stemming from genome-genome interactions of specific ant-fungus combinations (so-called inter-genomic epistasis; Wolf et al. 2000; Heath 2010; Wade 2007).

The range of *A. texana* is thought to be limited by cold temperatures in north Texas, low precipitation in west Texas, and a shallow water table to the east along the Mississippi Valley in Louisiana (Mueller et al. 2011b). Leafcutter ants can protect their fungal gardens against some temperature fluctuations (Mueller et al. 2011a), but at their northern range limits the warmest soil temperatures in winter (≈15°C) occur at depths below 5-10 meters, whereas more shallow depths, where gardens are maintained, are much colder (5-15°C) in winter. Consequently, fungiculture in leafcutter populations at their northern range limits must operate throughout winter at temperatures that would critically compromise survivorship of *L. gongylophorus* associated with tropical leafcutter ant species to the south. In an experiment testing for cold tolerance and desiccation resistance in the *L. gongylophorus* fungus, cold-adapted strains occur at northern sites across the range of *A. texana*, and cold-susceptible *L. gongylophorus* tend to occur at warmer southern sites in the Rio Grande Valley at the US-Mexico border (Mueller et al. 2011a). In contrast, there were no regional differences in desiccation resistance among strains (Mueller et al. 2011a).

### 2.2 Sample collection and processing

For genotype-by-sequencing using ddRADseq (Peterson et al. 2012), we used *A. texana* workers from 111 nests collected across Texas and Louisiana in 2000-2014 (Table S1; Mueller et al. 2011b). Mesosomas from three large workers per nest were washed three times in 100% ethanol to clean the integument, crushed with a sterile pestle in liquid nitrogen, then extracted with the Qiagen DNAEasy kit. *L. gongylophorus* samples from these nests had previously been genotyped with 12 microsatellite loci as described in Mueller et al. (2011b).

We digested 240-320 ng of *A. texana* DNA with NlaIII and EcoRI-HF (NEB) and prepared the ddRAD libraries following Peterson et al. (2012) using ‘flex’ barcoded adaptors. We used one male to generate a draft reference genome, which was assembled using ABYSS (Simpson et al. 2009). Methods for constructing ddRAD libraries and the reference genome are available in the electronic Supporting Information. Libraries were sequenced with Illumina Hiseq 4000 (ddRAD; 2 x 150bp) and Illumina Hiseq 2500 (draft genome; 2 x 100bp). The RAD-sequence information is available in NCBI BioProject PRJNA395768, and the draft genome of *A. texana* is available at NCBI as accession QEPB00000000.

### 2.3 Bioinformatics

We used Stacks 1.40 (Catchen et al. 2011) to demultiplex the ddRAD sequences and aligned reads to the *A. texana* genome with BWA mem v0.7.12 (Li 2013), retaining only reads that mapped in a perfect pair. We genotyped samples using samtools mpileup/bcftools v1.3.1 (Li et al. 2009), specifying a MAPQ score >20, maximum read depth of 500, mapping quality adjusted to 50, BAQ disabled, and the consensus calling model. We removed sites with individual read depths less than 15, genotyping quality less than 20, and sites present in less than 80% of individuals using vcftools 0.1.15 (Danecek et al. 2011). We used only single-nucleotide polymorphisms (SNPs) from bi-allelic loci in analyses. SNPs were thinned to a maximum of one per 100 kb to remove linked sites.

### 2.4 Spatial principal component analysis (sPCA) of population structure

SPCA is a model-free method that incorporates spatial relationships among samples into a principal components analysis to infer genetic structure among individuals or populations (Jombart et al. 2008). To implement sPCA, we randomly chose one ant per nest and specified a Gabriel graph to define spatial connectivity (Figure S2a) using the adegenet package (Jombart & Ahmed 2011) in R v3.2.1. We visually inspected screeplots of the eigenvalues to determine how many sPCAs to retain for analysis. To accommodate the apparent variation in ploidy among *L. gongylophorus* individuals (Carlson et al. 2017), we set the ploidy level for each individual as the maximum number of microsatellite alleles counted across any of the 12 loci. Alleles were coded as dominant markers as recommended for polyploids by Falush et al. (2007). To assess whether global and local structures were present in our data, we implemented the global.rtest and local.retest functions in the adegenet package (Jombart & Ahmed 2011). We visualized the results by plotting the sPCA scores onto a map of Texas and Louisiana using ggmap (Kahle & Wickham 2013).

### 2.5 Genetic diversity

We assessed the relationship between genetic diversity and geography in the ant *A. texana* by regressing heterozygosity on longitude and latitude. *L. gongylophorus* fungi exhibits inter-individual variation in ploidy, so we used allele richness as a measure of genetic diversity. Allele richness is highly correlated with heterozygosity for microsatellite markers (e.g., the review by Eckert et al. 2008 calculated a correlation of r=0.81 for estimates from 15 published studies), and thus serves as an acceptable proxy for genetic diversity. We generated subsets of *L. gongylophorus* samples grouped by their estimated ploidy level [see *Spatial principal component analysis (sPCA) of population structure*] to prevent confounding comparisons of allele richness among individuals that could be solely due to ploidy. We used Spearman rank correlation to compare allele richness with longitude and latitude because allele richness could not be transformed to meet the assumptions of linear regression. We visualized results using the R package ggmap (Kahle & Wickham 2013).

### 2.6 Cluster-based analysis of population structure

We assessed population-genetic structure in *A. texana* using ADMIXTURE (Alexander et al. 2009) to complement the sPCA. Unlike sPCA, ADMIXTURE clusters samples populations using a maximum-likelihood-based population-genetic model that assesses the contribution of K ancestral populations to each individual genome. To determine the K for the analysis, we applied ADMIXTURE’s implementation of cross-validation (n=5 folds, K=1-5) and selected K from the model with the lowest error. Admixture proportions were then plotted using the maps package in R (Becker et al. 2016) to visualize genetic structure across the range of *A. texana*.

### 2.7 Contributions of cultivar genotype and climate to genetic variation in *Atta texana*

We used BEDASSLE (Bradburd et al. 2013) to assess the relationship between ecological factors (temperature, precipitation, and fungal genotypic cluster) and genetic differentiation among *A. texana* nests. We downloaded mean temperature and precipitation for the years 1998-2010 from the National Centers for Environmental Information and assigned the fungal genotype cluster for each nest as inferred already in an earlier microsatellite-marker analysis (a total of 79 variable markers across 12 highly-polymorphic microsatellite loci; Mueller et al. 2011b). We initiated two replicate Markov Chain Monte Carlo (MCMC) runs of the overdispersion model using the BEDASSLE package in R v3.2.1. Each run used a different random seed value to ensure independence, and included ~10 million MCMC iterations after burn-in (the first 20% of the run), with parameter values sampled every 5000 generations. Model convergence was evaluated using graphical functions implemented in the BEDASSLE package. We also evaluated model fit by comparing naturally observed Fst values with 100 posterior predictions drawn randomly from the MCMC iterations. The effect size of each ecological parameter (αE) relative to isolation by distance was then calculated by dividing their posterior distributions by the posterior distribution of the effect size of geographic distance (αD) on genetic differentiation; this step permits the simultaneous estimation of the effect sizes of all parameters using the same units and relative to change in population-structure due to geographic distance, as explained in Bradburd et al. (2013) and Weber et al. (2017). Highest posterior density credible intervals for each effect were calculated using the coda package in R (Plummer et al. 2006). Finally, we calculated mean posterior estimates for within-population allelic correlations (□), and then converted these to F-statistics (analogous to inbreeding coefficients) following Bradburd et al. (2013). We used a linear model to test for an association between fungal type and F-value.

## 3 Results

Using genotyping-by-sequencing ddRADseq methods, we identified 4003 single-nucleotide polymorphisms (SNPs) in *A. texana* leafcutter ants collected from 111 nests covering the entire range of this ant species. A screeplot of the eigenvalues in the ant sPCA analysis indicated three positive eigenvalues (representing global structure) for retention (Figure S2b). Plotting the spatial and variance components of these eigenvalues reinforced that these three axes were distinguishable from the rest of the eigenvalues, and thus suitable for interpretation (Figure S2c) (Jombart & Ahmed 2011). Negative eigenvalues (representing local structure) were relatively small (Figure S2b), suggesting weak local structure in the data. Formal tests using MCMC simulation confirmed that genetic variation was significantly associated with global spatial structure (p<0.001) but not with local spatial structure (p=0.99), and we therefore analyzed only global structures further.

The fungus microsatellite-marker dataset (Mueller et al. 2011b) of *Leucocoprinus gongylophorus* fungi cultivated by the same 111 nests of *A. texana* included 79 variable markers distributed across 12 loci (2-10 alleles per locus for this multinucleate, polyploid fungus). We identified one positive eigenvalue for retention in the analysis (Figure S3a,b), and as in the ant dataset, little evidence of local spatial structure in the screeplots (Figure S3a). MCMC simulation confirmed that genetic variation was significantly associated with global (p=0.003) but not local structures (p=0.49). We thus only focused on global structures in the remaining analyses.

### 3.1 Genetic structure of *Atta texana*

Spatial genetic structure among nests was strikingly different between the ants and their fungi. The first sPCA axis in the ants revealed a broad cline spanning from the Rio Grande at the US-Mexico border (older populations of the range expansion) to Louisiana (Figure 1a). This cline likely results from a strong effect of isolation-by-distance (Figure S4) on genetic variation in the ants. A concurrent decline in heterozygosity was also evident from the Rio Grande to Louisiana (Figure 2A). We found a negative relationship between heterozygosity (range: 0.0005 - 0.26) and latitude [β (95% CI) = −0.011 (−0.014, −0.008), R^2^=0.33], and between heterozygosity and longitude [β (95% CI) = −0.007 (−0.010, −0.003), R^2^=0.11]. Together, these results indicate that genetic drift reduced genetic variation in the ants as they expanded their range northeast, and gene flow had insufficient time to erode the genetic signature of this range expansion.

**FIGURE 1.**
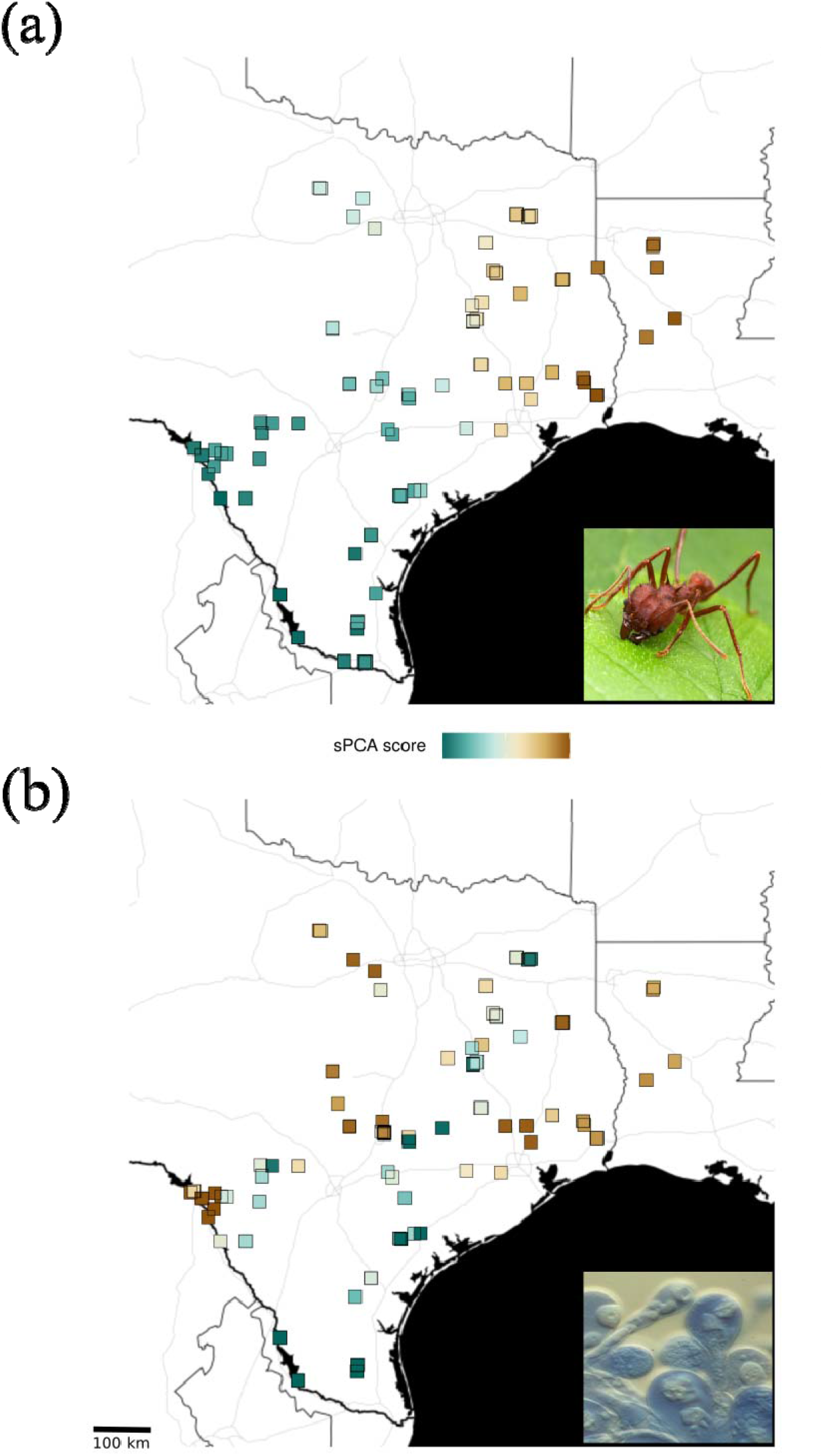
Global scores from the first axis of the spatial principal component analysis of (a) the leafcutter ant *Atta texana* and (b) *Leucocoprinus gongylophorus* fungal cultivars collected from the same nests. Inset in (a) is an *A. texana* worker (photo courtesy of Alex Wild), and inset in (b) gongylidia (hyphal-tip swellings) of *L. gongylophorus* cultivar (photo by Ulrich Mueller).

**FIGURE 2.**
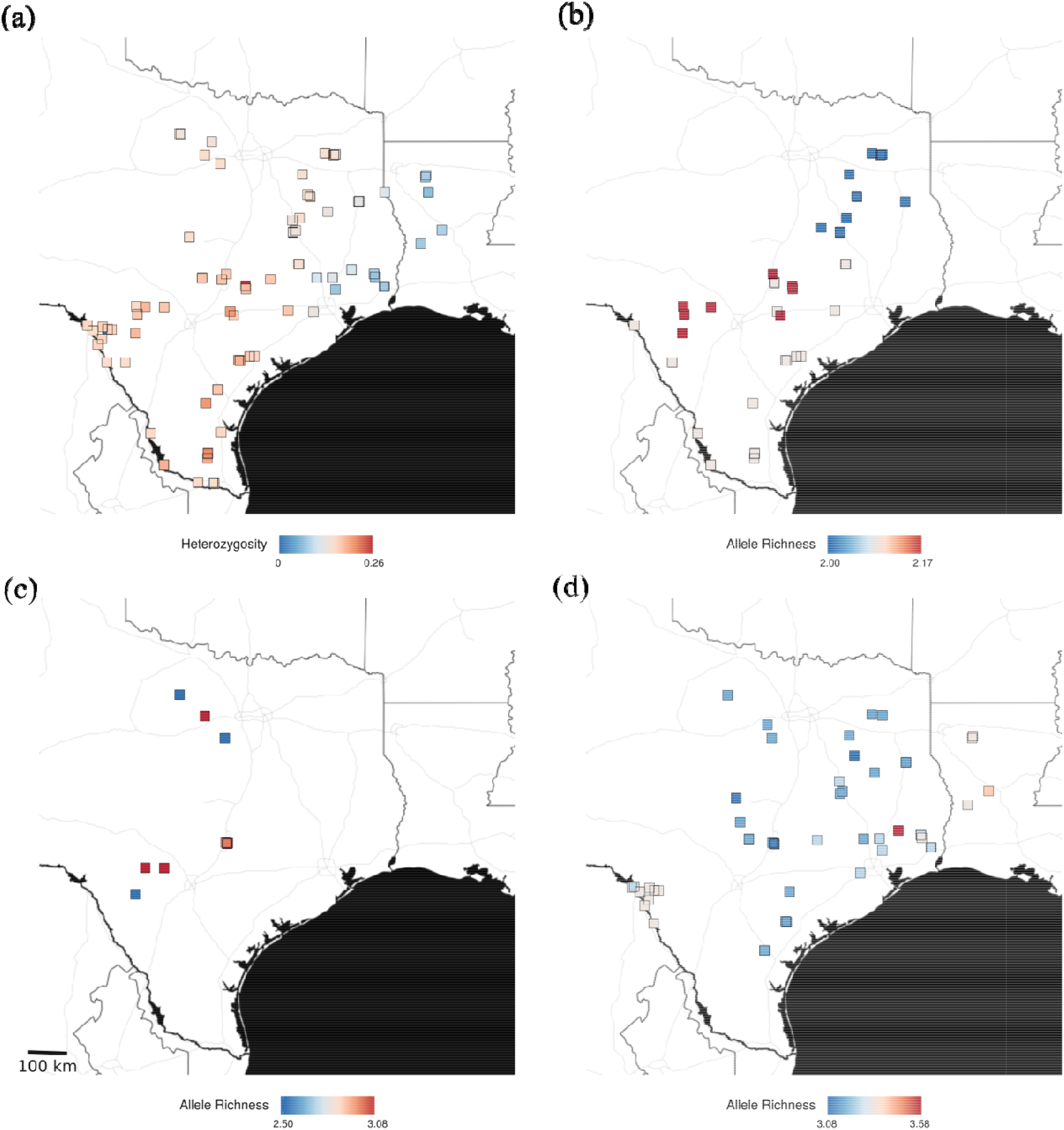
Genetic diversity in (a) the leafcutter ant *Atta texana* and (b) putative triploid, (c) putative tetraploid, and (d) putative pentaploid *Leucocoprinus gongylophorus* fungal cultivars.

The second and third sPCA axes revealed additional genetic structure. The second sPCA axis differentiated a cluster of nests along a corridor from north Texas to eastern Texas (Figure S5a), the third sPCA axis differentiated a cluster of nests in central Texas (Figure S5b).

ADMIXTURE, which relies on an evolutionary model to assign admixture proportions, revealed similar results as the sPCA. The crossfold validation estimated K=3 as the most likely number of clusters in the data (Figure S6a). The first cluster is comprised of nests in southwest and central Texas, the second of eastern and north Texas, and the third of eastern Texas/Louisiana (Figure S6b). Qualitatively, these divisions correspond to the patterns revealed in the first sPCA axis, which accounts for most of the variation in the data (Figure S2b,c).

### 3.2 Genetic structure of *L. gongylophorus* fungus

Spatial PCA of the fungi revealed a patchwork of genetically similar cultivars distributed across the range (Figure 1b). This is in agreement with (Mueller et al. 2011b) documenting a similar pattern when analyzing the same dataset using a population-genetic model-based approach (STRUCTURE). Allele richness decreased with longitude and latitude in putative triploid samples (longitude: Spearman’s rho = −0.70, p < 0.001; latitude: rho = −0.53, p < 0.001, n = 48; Figure 2b), but not in putative tetraploids (longitude: rho = −.35, p= 0.3; latitude: rho = −0.26, p=0.5, n = 10; Figure 2c) or putative pentaploids (longitude: rho = 0.14, p=0.3; latitude: rho = - 0.29, p = 0.1, n=60; Figure 2d). The decline in genetic diversity in triploid samples was due to a cluster of genetically identical genotypes with low allelic richness in northeast Texas (Figure 2b).

### 3.3 Contributions of fungal genotype and climate to genetic variation in *Atta texana*

We found that temperature, precipitation, and fungal genotype-cluster (so-called M- or T-fungi) were all associated with genetic differentiation in *A. texana* (Table 1). One degree Celsius increase in temperature was equivalent to 60.2 km (mean of two MCMC runs) of genetic differentiation due to isolation-by-distance (IBD) among nests, while every inch increase in precipitation was equivalent to 52.3 km of genetic differentiation due to IBD.

**Table 1.** Medians (credible intervals) of posterior effect sizes of geographic distance (αD) and ecological factors (αE) on genetic differentiation in the ant *Atta texana*, listing output from two independent MCMC runs on separate lines in each cell.

| Source | $\alpha$ (95% Credible Interval) | $\alpha_E / \alpha_D$ |
| --- | --- | --- |
| **Geographic distance** | 1.34e-06 (8.17e-07, 1.89e-06)<br>1.52e-06 (9.14e-07, 2.18e-06) |  |
| **Temperature**<br>**(per °C)** | 7.78e-05 (5.85e-09, 2.99e-04)<br>8.86e-05 (3.00e-08, 3.78e-04) | 59.1 (4.33e-03, 211)<br>61.4 (1.75e-02, 248) |
| **Precipitation**<br>**(per inch annual rainfall)** | 6.87e-05 (4.12e-05, 9.86e-05)<br>7.97e-05 (4.56e-05, 1.14e-04) | 51.5 (38.9, 68.1)<br>53.0 (40.3, 68.5) |
| **Fungal genotype cluster** | 8.64e-04 (3.52e-04, 1.62e-03)<br>1.12e-03 (4.60e-04, 1.95e-03) | 673 (271, 1210)<br>766 (373, 1240) |

Genetic differentiation between ants associated with either M- vs T-fungi was relatively large, comparable to 719 km of IBD. Relative to the abiotic factors, this is equal to an 11.9 C° difference in temperature (719 km / 60.2 C°/km), or a 13.7 inch difference in precipitation (719 / 52.3 in/km) (Figure 3). Fungal genotype-cluster thus had a much larger effect than temperature, which differs by 7.1 C° average January temperature (interquartile range = 2.7 C°) from the coldest to the warmest locations, but less than that of precipitation, which differs by 86.7 inches annual rainfall (interquartile range = 60.2 inch) between the driest and wettest locations. Our BEDASSLE models also produced highly variable F-statistics across ant populations (range = 3.96e-4 to 0.528, SD = 0.133), with high values resulting from either high inbreeding or poor model fit. However, there was no association between F-values and fungal type (linear model: t *=* 0.26, p = 0.80).

**FIGURE 3.**
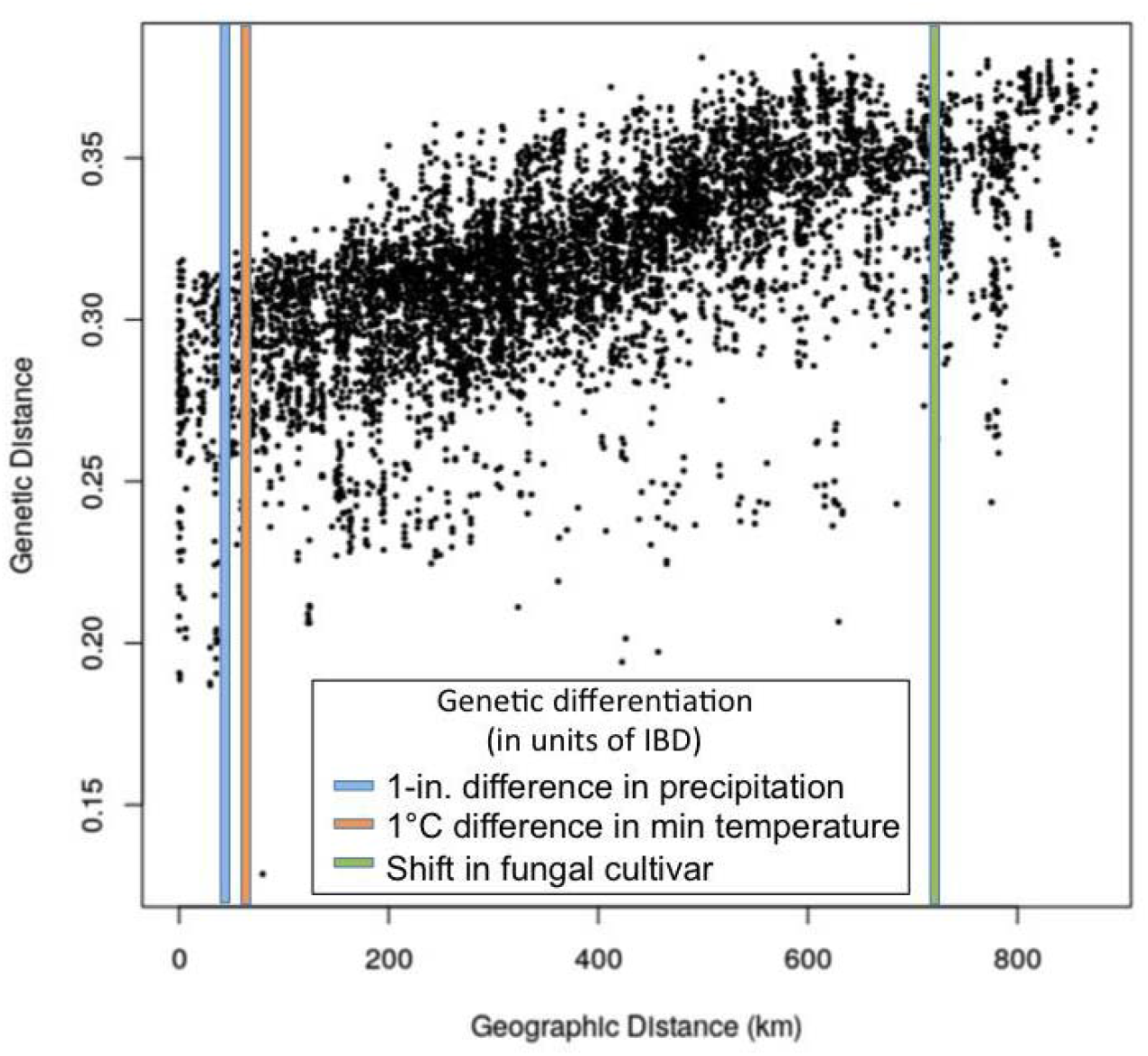
Isolation-by-distance (IBD) and isolation-by-ecology in the leafcutter ant *Atta texana*. Genetic distance is Edward’s angular distance (method 2 in the dist.genpop function of the adegenet R package). Colored bars indicate the impact of ecological differences relative to IBD (see Table 1 for additional information).

## 4 Discussion

The ecological and evolutionary processes driving and limiting range expansions have been debated since Darwin suggested that both abiotic and biotic factors can limit species boundaries (Darwin 1859; Sexton et al. 2009; Schoville et al. 2012). The role of mutualisms in facilitating or inhibiting range expansions, however, has received far less attention than other species interactions, such as competition and predation (Gilman et al. 2010; Lavergne et al. 2010; Urban 2011). We found that the range expansion during the last 11,000 years by the northernmost leafcutter ant *Atta texana* across the south-central USA (Texas and Louisiana) left a strikingly different genetic footprint on each partner in this obligate ant-fungus mutualism, indicating that the evolutionary processes accompanying expansions can be markedly different in obligately interdependent species. Second, we observed that genetic differentiation among the ant hosts depends upon the genotype of their symbiotic fungal partner (T- or M-fungi), and that the strength of this effect was stronger over local geographic scales than either of two abiotic environmental factors, temperature and precipitation. Our results support theory predicting that (a) range expansion can leave an indelible genetic signature (Figures 1a and 2a); and (b) both symbiotic associations and environmental factors impact genetic differentiation of a host (Figure 3).

### 4.1 Magnitude of the effects of symbiont type, temperature, and precipitation on genetic differentiation in the ant *Atta texana*

#### 4.1.1 Symbiont type

Isolation-by-environment occurs when genetic differences among populations increase along an environmental gradient independent of their geographic distance (Wang & Bradburd 2014). We found that temperature, precipitation, and fungal genotype-cluster were all associated with genetic differentiation across the range of *A. texana*, suggesting that both abiotic factors and ant-fungus interactions influenced evolution during range expansion. Genetic differences between *A. texana* associated with M- or T-fungi of *L. gongylophorus* were large relative to the abiotic factors (Figure 3), equivalent to the amount of isolation-by-distance of populations 719 km apart (e.g., between populations at the tip of south Texas to those in central Texas, about half the species range). Three hypotheses can explain these results:

First, genetic interactions between *A. texana* and *L. gongylophorus* could affect survival and reproduction, resulting in correlations between genetic variation in ant and fungal genotypes because specific ant-fungus combinations are selectively favored. Genetic interactions (i.e., epistasis) are traditionally studied between nuclear genes, but epistasis can also occur between genomes (Wolf et al. 2000; Gilbert 2002; Wade 2007), such as nuclear-mitochondrial (Dowling et al. 2008; Ballard & Melvin 2010), nuclear-chloroplast (Wolf et al. 2000), or host-symbiont interactions (Mueller et al. 2005; Wade 2007; Heath 2010). In leafcutter ants, novel ant-fungus combinations might arise by mutation and hybridization in the fungi, or cultivar switching of ant colonies between T- and M-fungi. We expect beneficial, co-adapted genome-genome combinations will be more likely to co-propagate than mismatched (i.e., selectively inferior) combinations. Genome-by-genome fitness effects of mutualisms have been tested in controlled common-garden experiments, for example in endophytic or rhizobial mutualists of plants (Gilbert et al. 2010; Heath 2010), whereas our survey suggests that inter-genomic epistasis for mutualisms could operate under natural conditions across the range of a host species.

Although less plausible than inter-genomic epistasis, there are two alternative explanations for fungus-associated genetic differentiation. First, genetic differentiation between ants associated with either M- or T-fungi could be due to a spurious correlation with another factor, such as if ants with different fungal types evolved in allopatry. Ant nests cultivating either M- and T-fungi, however, are sympatric across much of the range, occurring as close as 80 meters apart in some instances (Mueller et al. 2011b). With no obvious physical or environmental barriers preventing cultivar switching between nests and with ample opportunity for gene flow between ants with different fungal types, genetic differentiation of ants with M- and T-fungi due to allopatry therefore is an unlikely explanation.

As a second alternate explanation, the association of genetic variation between ants and fungal genotype (M- or T-fungi) could be due to a demographic process unrelated to ant-byfungal interactions *per se*, such as population growth of specific ant lineages cultivating either M- or T-fungi. Because cultivars are vertically transmitted, fungal strains could ‘hitchhike’ with a successful ant lineage much like loci in linkage disequilibrium with selected genes during a selective sweep (Vitti et al. 2013). The most likely scenario for this would be a ‘double invasion’ of *A. texana* expansion into Texas, one invasion by ants bearing the T-fungus and a second bearing the M-fungus. Because gene flow among ant lineages would erode genetic differences between M- and T-cultivating ant lineages, the second invasion would have to occur quickly relative to the rate of interbreeding between ant lineages. However, given the slow dispersal rate of *A. texana* (see below) and sympatry between M- and T-cultivating ants across most of the range of *A. texana* (Mueller et al. 2011b), this scenario seems very unlikely. Moreover, the lack of association between fungal type and F-values in our BEDASSLE models also suggest that a purely demographic explanation is unlikely. Models reconstructing different possible demographic histories (Schraiber & Akey 2015) are needed to formally test this hypothesis.

#### 4.1.2 Temperature and precipitation

In addition to fungal genotypic cluster, temperature and precipitation were also associated with genetic differentiation in *A. texana*, independent of isolation-by-distance in the ants. Genetic differentiation increases more rapidly with temperature in *A. texana*, however the absolute potential for differentiation was greater along the east-west precipitation gradient because of the wider cline in rainfall (i.e., annual precipitation increases 210% from West Texas to Louisiana, while annual temperature increases 42% from south Texas to north Texas; Figure S1). Wang & Bradburd (2014) identified several processes that generate genome-wide associations between genetic differentiation and environment across the landscape: (i) local adaptation and selection against maladapted immigrants; (ii) biased dispersal of genotypes to a preferred habitat, leading to a correlation between genotype and habitat regardless of whether selection acts once migrants arrive; and (iii) isolation-by-distance, a common confounding factor in landscape genomics that arises because environmental differences are often also greatest among geographically distant samples.

Biased dispersal is unlikely to account for our results because the spatial scale of our sampling over the environmental gradient far exceeds the dispersal distance of female leafcutter ants during the mating flight. In other words, while queens can choose specific microhabitats for nest building (e.g., shaded areas under trees), they would need to fly much further than they can to sample habitats that differ significantly in mean annual temperature and precipitation. To illustrate, the invasive leafcutter ant *Acromyrmex octospinosus* expanded across the Caribbean island of Guadeloupe about ~0.5 km per year, generating isolation by distance at geographic scales as small as 1000 km^2^ (Mikheyev 2008), less than 1% of the area examined here for *A. texana*. Dispersal distance of *A. texana* females in the field is not known, but dispersal is estimated to be less than 10 km per mating flight (Moser 1967) or much less (Mueller et al. 2011b). Wang & Bradburd’s (2014) third factor, isolation by distance, can also lead to spurious correlations between genetic differentiation and environmental factors, however isolation by distance was controlled for in our analysis. Local adaptation and selection against maladapted immigrants thus remains a viable explanation for the observed genome-wide differentiation over the cline in temperature and precipitation.

Temperature and precipitation have wide-ranging effects on physiological function of insects, and leafcutter ants have evolved adaptations to cope with cold, heat, and drought (Bollazzi & Roces 2002, 2010a,b; Bollazzi et al. 2008; Ruchty et al. 2010). *A. texana* exhibits behavioral adaptations to provide environments suitable to maintain the productivity of fungal cultivars, for example by building garden chambers at soil depths that meet an acceptable range of temperature and humidity, or by moving fungal gardens to depths with optimal environmental conditions (Mueller et al. 2011a). The observation that fungal cultivars from colder locales exhibit greater cold tolerance suggest that environmental gradients in temperature are steep enough to have led to a response to selection (Mueller et al. 2011a), at least in the fungus.

An important caveat is that the observed genome-wide differentiation can be caused by other, unknown factors correlated with the environmental gradients. While BEDASSLE explicitly controls for the effect of isolation by distance, we cannot rule out that unmeasured variables other than temperature and precipitation might be ultimately responsible to the observed patterns. One such factor could be the SW-NE direction of the range expansion, and therefore genetic drift, which is not completely orthogonal to the gradients in temperature (north/south) and precipitation (east/west) (Figure S1). Nevertheless, our results are consistent with parallel evolution in both *A. texana* ants and associated *L. gongylophorus* fungi along the temperature gradient (Mueller et al. 2011a), and evolution of *A. texana* but not the fungi in response to the precipitation gradient (Figure S1). Laboratory experiments testing the performance of ants under cold and desiccation stress, as well as genomic analyses of selection, are required to further substantiate these adaptive hypotheses.

### 4.2 Spatial associations between genetic variation in ants and their fungal cultivars

#### 4.2.1 Population genetic structure in the leafcutter ant *Atta texana*

We found (i) a broad cline in genetic differentiation (Figure 1a) and (ii) a reduction in heterozygosity (Figure 2a) across the entire range of *A. texana*, from the Rio Grande at the US-Mexico border to northeast Texas and Louisiana. These observations are consistent with a model of a north-eastward expansion of *A. texana* following the glacial retreat in the Pleistocene, accompanied by limited gene flow across the range as new habitat was colonized by the ants. The continuous and graded cline in genetic differentiation is likely due to the lack of prominent physical barriers to dispersal across the range of *A. texana* (i.e., there are no mountain ranges or large river-basins that prohibit the movement of female reproductives during dispersal flights). Instead, gene flow is likely restricted by the short dispersal distance of female reproductives during their annual mating flight (less than 10km, or far less; Moser 1967; Mueller et al. 2011b), generating strong signals of isolation-by-distance (Figure S4) and clinal population structure (Figure 1a).

The observed decline in heterozygosity in *A. texana* towards the range limit (Figure 2a) is expected if genetic variation existing in ancestral populations was eroded by drift as founding queens dispersed along the front of the expanding population (Eckert et al. 2008; Excoffier et al. 2009; Sexton et al. 2009). This cline in population structure and loss of genetic diversity across the range might have important evolutionary consequences because (i) smaller effective population sizes at the wave front increase drift, weakening the efficacy of selection; (ii) deleterious mutations can ‘surf’ to high frequency along the wave front due to genetic drift; and (iii) immigration of alleles from the center of the range can constrain the speed of adaptation to conditions at the range edge (Mayr 1963; Nei et al. 1975; Kirkpatrick & Barton 1997; Klopfstein et al. 2005; Excoffier & Ray 2008; Sexton et al. 2009, 2014; Peischl et al. 2013). These factors are likely compounded in mutualisms if both partners in the symbiosis must adapt independently to conditions at the wave front to survive, and when a symbiont propagates asexually, as is the case for the largely asexual fungi cultivated by *A. texana* (Mueller et al. 2010; Mueller et al. 2011b; Marti et al. 2015). Genetic drift operating at the expanding range front might also reduce the number of possible combinations of ant and fungal genomes co-existing in a population at the range edge, limiting the opportunity for advantageous genome-genome interactions (i.e., coadaptations) that facilitate adaptation.

#### 4.2.2 Population-genetic structure in the fungal cultivar compared to ant host

The striking contrast in the spatial distribution of genetic variation of *L. gongylophorus* fungi compared to the ant hosts suggests that the evolutionary forces operating on each partner in this mutualism can be highly discordant. In contrast to the smooth cline in population structure (Figure 1a) and the spatially contiguous corridors of shared genetic diversity in the ant host (Figures S5a & S5b), population structure in the fungi is patchily distributed across the range (Figure 1b) (Mueller et al. 2011b), and there was no clear reduction in genetic diversity of the fungal symbionts in the direction of the host’s population expansion (Figures 2b-d). One possible explanation for this mismatch is that gene flow might occur over larger spatial scales in *L. gongylophorus* fungi than in the ants. While *L. gongylophorus* fungi are only known to occur within leafcutter nests and are vertically propagated by dispersing females during the mating flight, fungal cultivars are also occasionally horizontally exchanged among nests (Mikheyev et al. 2007; Mikheyev et al. 2010; Mueller et al. 2017; Mueller et al. 2018). Specifically, *L. gongylophorus* genotypes are clustered at local geographic scales, consistent with vertical transmission, but identical cultivar clones can also be shared between nests over large distances (Mueller et al. 2011b; Mueller et al. 2017). In contrast to these evolutionary forces affecting the cultivated fungi, genetic drift and limited long-distance gene flow appears to be more important in the evolutionary dynamics in the ants (Figure 1a).

## 5 Conclusion

The observed differences between *A. texana* and *L. gongylophorus* in the importance of adaptation, genetic drift, and gene flow has implications for evolution during the northward range expansion of leafcutter ants since the end of the last glaciation 11,000 years ago. Average temperature in January at the range front in northern Texas (−3 to 3 C°), for example, is currently about 10 C° lower than in the south (7 to 10 C°) (Mueller et al. 2011a), and fungi cultivated by *A. texana* in northern populations are more cold-tolerant than those in southern populations (Mueller et al. 2011a). If north-south gene flow in *L. gongylophorus* has occurred, it has not precluded local adaptation of the fungus to temperature. Whether local adaptation at the range front has similarly occurred in the ant hosts has yet to be tested experimentally, but the limited opportunity for dispersal by the ants does make it unlikely that gene flow precludes range-limit adaptation in the ants. Genetic drift in *A. texana*, on the other hand, reduces the number of possible combinations of interacting alleles between ants and their fungal cultivars, possibly limiting the opportunity for beneficial inter-genomic synergisms that facilitate adaptation at the range front. Whether range expansion of leafcutter ants is impacted by interspecific interactions therefore depends on complex interactions between selection, gene flow, and drift acting in parallel in both hosts and symbionts to either maintain or erode adaptive inter-genomic synergisms.

## Acknowledgements

We thank numerous landowners for permission to collect on their properties, Gideon Bradburd for advice on the BEDASSLE analysis, Mark Kirkpatrick for advice on population-genetic interpretations, Dan Bolnick for material support, and Alex Wild for the ant photo. The work was funded by National Science Foundation award DEB-1354666 and the W.M. Wheeler Lost Pines Endowment from the University of Texas at Austin.

## Data Accessibility

RAD-sequence data are available in NCBI BioProject PRJNA395768, and corresponding sampling metadata for *Atta texana* collections are available as a tsv file in the Supporting Information. The draft genome of *Atta texana* is available in NCBI accession QEPB00000000.

## Author Contributions

C.C.S., J.N.W. U.G.M., F.R., M.B., K.K., and J.N.S. planned sampling and coordinated research. C.C.S. and J.N.W. designed the sequencing strategy and conducted population-genomic analyses. C.C.S. archived data. A.S.M. assembled the draft genome of *Atta texana*. C.C.S., U.G.M., and J.N.W. wrote the manuscript. U.G.M., K.K., M.B., F.R., and J.N.S. procured funding. All authors revised the manuscript and approved the final manuscript.

## Orcid

Ulrich G. Mueller http://orcid.org/0000-0003-2677-8323 Alexander S. Mikheyev http://orcid.org/0000-0003-4369-1019

## SUPPORTING INFORMATION

### Landscape genomics of an obligate mutualism: discordant population structures between a leafcutter-ant and its fungal cultivars

Smith, C.C., J.N. Weber, A.S. Mikheyev, F. Roces, M. Bollazzi, K. Kellner, J. Seal, U.G. Mueller

### Genomic Methods

For genotyping-by-sequencing, we chose 111 *A. texana* nests collected across Texas and Louisiana in 2000-2014 (Table S1; Mueller et al. 2011a,b). Mesosoma from three large *A. texana* workers per nest were retrieved from collections stored in 100% ethanol at −80C°, washed three times in 100% ethanol to clean the integument, crushed with a sterile pestle in liquid nitrogen, then extracted with the Qiagen DNAEasy kit. DNA concentration was assessed fluorometrically using Quant-iT TM Picogreen dsDNA Assays (Life Technologies) and an Infinite Pro M200 Pro microplate reader (TECAN), resulting in 8.1 ng/μl + 5.3 (mean + SD) of DNA per sample for library preparation. To prepare the ddRAD libraries, we digested 240-320 ng of DNA with 5 units of NlaIII and 5 units of EcoRI-HF (NEB) in 80 μl reactions overnight, normalized the resulting product to 90 ng after cleanup with Sera-Mag™ Magnetic SpeedBeads™ (GE Healthcare, see protocol in Rohland & Reich 2012) and ligated the DNA with a 5x excess of ‘flex’ barcoded adaptors (0.16 pg of P1, 0.14 pg of biotin-labeled P2) using 400U of T4 ligase (NEB) in a 40 μl reaction. Barcoded products were pooled, cleaned, and 426-451 bp fragments were selected using a PippenPrep (2% agarose gel; Sage Sciences). We then applied 4 μl of M-270 Streptavidin-coated Dynabeads (Life Technologies) per pool to purify biotin-labeled fragments. Illumina-compatible adaptors were added using 12 cycles of PCR with Phusion Hi-Fidelity PCR Kit (NEB) on six replicates of each pool to mitigate “jackpot” effects during amplification. Library replicates were combined and analyzed on a 2100 BioAnalyzer (Agilent Technologies) for purity. Libraries were sequenced on an Illumina Hiseq 4000 (2 x 150bp PE) at the Genome Sequencing and Analysis Facility (GSAF) at the University of Texas at Austin.

We generated a draft reference genome (v 0.1) from one *Atta texana* male collected from the Brackenridge Field Lab (Austin, TX) at the University of Texas at Austin. We extracted the DNA as above and sent the sample to GSAF for library preparation and sequencing. Genomic DNA was sheared at GSAF to a 475-bp median fragment length using the Covaris S2 ultra-sonicator, and the library was prepared using the NEBNext DNA Library Prep Master Mix following the manufacturer’s instructions. Intermediate library fragments were purified after end repair and dA tailing steps using MinElute columns (Qiagen). The library was sequenced on an Illumina HiSeq 2500 using a paired-end 2X100 run that yielded 240 million reads. The *A. texana* genome was assembled in ABYSS (Simpson et al. 2009), with a range of kmer values from 25 to 95. We then chose the assembly with the longest N50 (7211 bp contigs and 10,786 bp scaffolds).

---

**TABLE S1.**Metadata for *Atta texana* are available as a tsv file in the Supporting Information.

**FIGURE S1.**
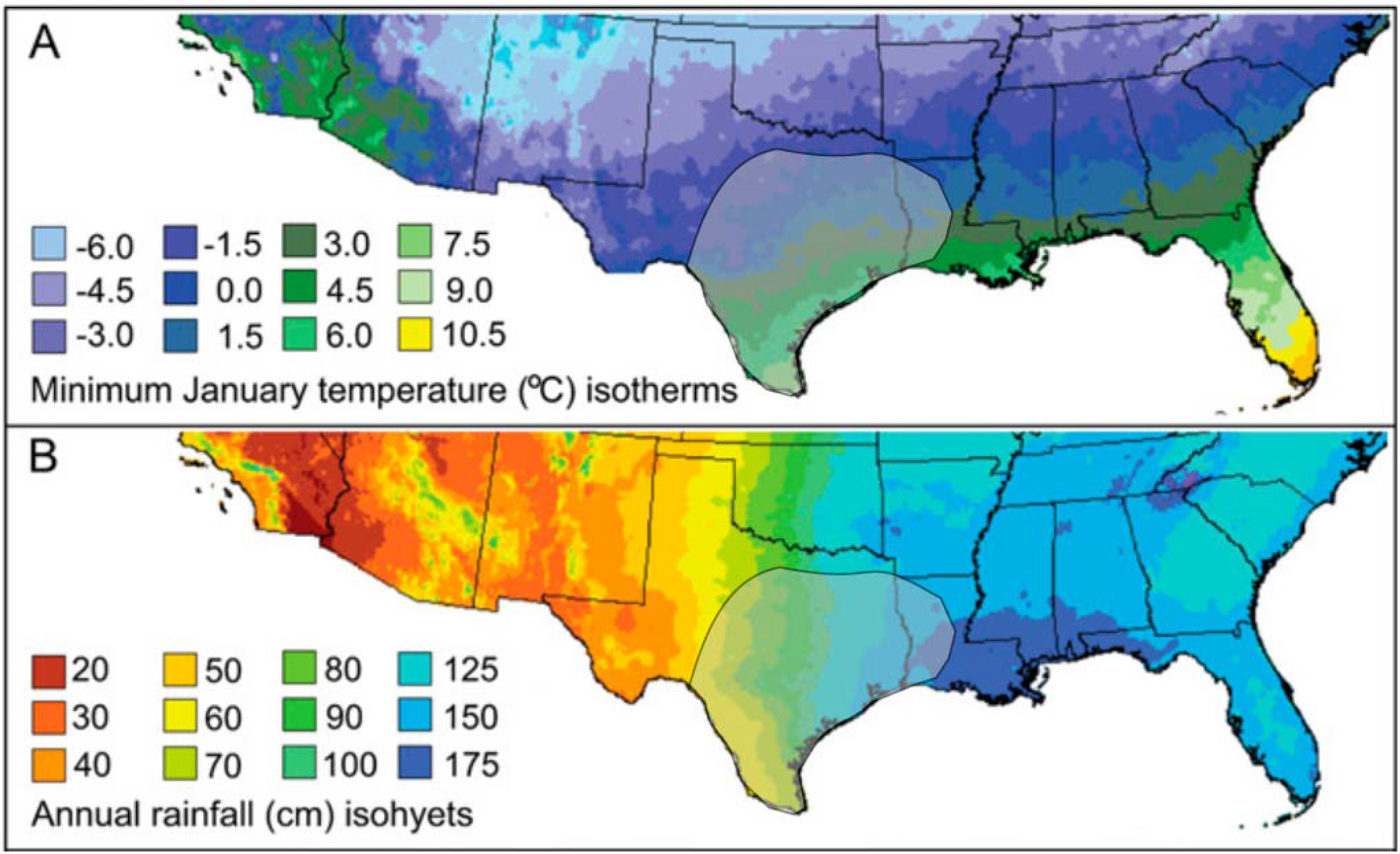
(A) Isotherm map of average minimum air temperature (in degree centigrade, °C) in January for the southern United States. (B) Isohyet map of average annual precipitation (in centimeter rainfall, cm). The approximate distribution of *A texana* in the USA is outlined in grey. The figure is modified from Mueller et al. (2011a).

**FIGURE S2.**
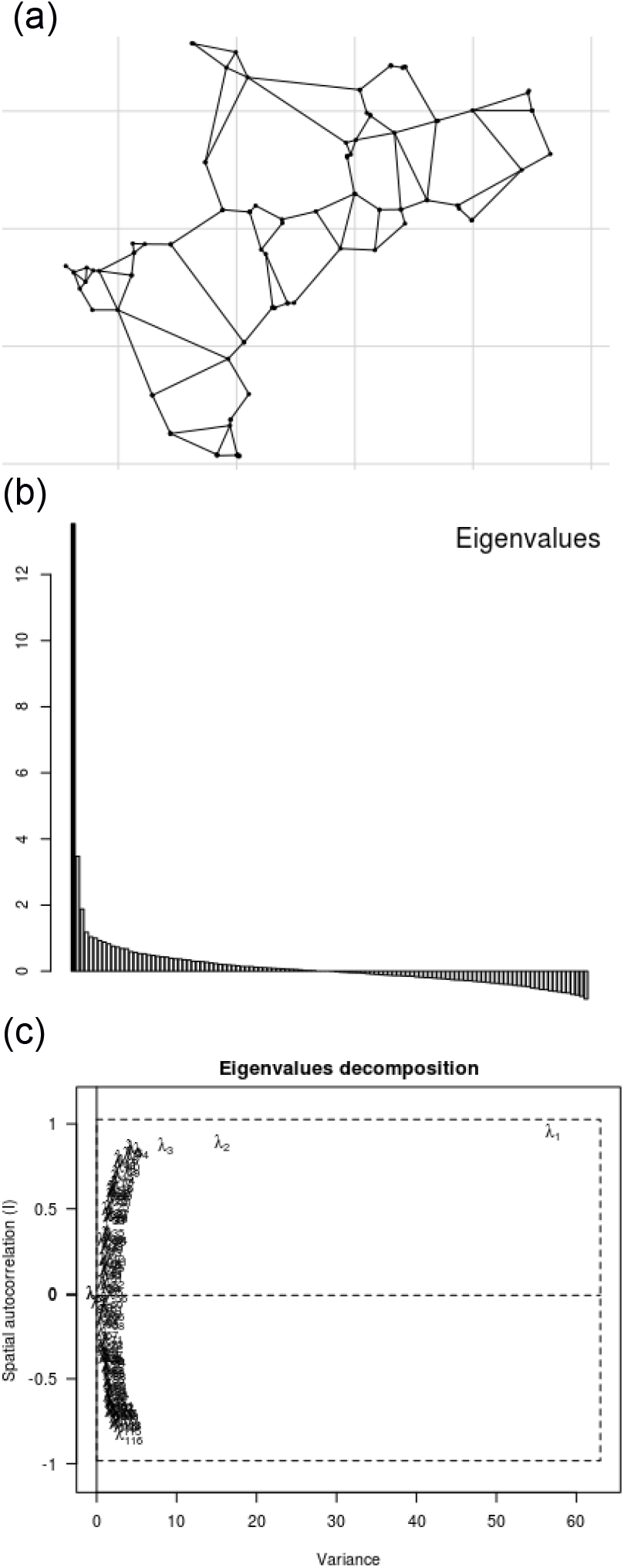
Spatial analysis of principal components for the ant *Atta texana*. (a) Nest connectivity, (b) screeplot of eigenvalues, and (c) relationship between spatial autocorrelation among nests and the total variance explained by each eigenvalue. Positive eigenvalues in (b) correspond to global structures and negative values correspond to local structures. Subscripts in (c) refer to the nth eigenvalue.

**FIGURE S3.**
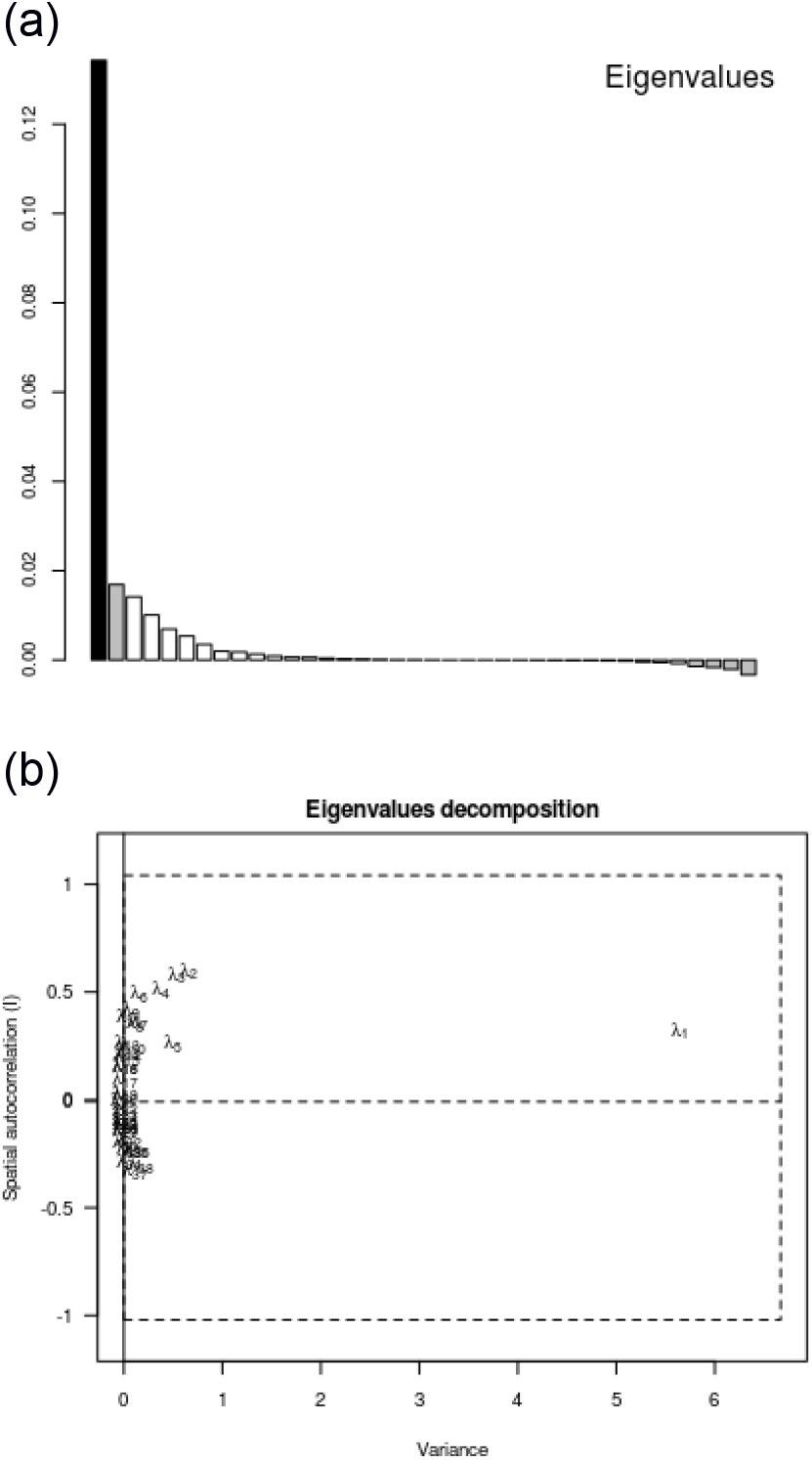
Spatial analysis of principal components for *Leucocoprinus gongylophorus*. (a) Screeplot of eigenvalues, and (b) relationship between spatial autocorrelation among samples and the total variance explained by each eigenvalue. Positive eigenvalues in (a) correspond to global structures and negative values correspond to local structures. Subscripts in (b) refer to the nth eigenvalue.

**FIGURE S4.**
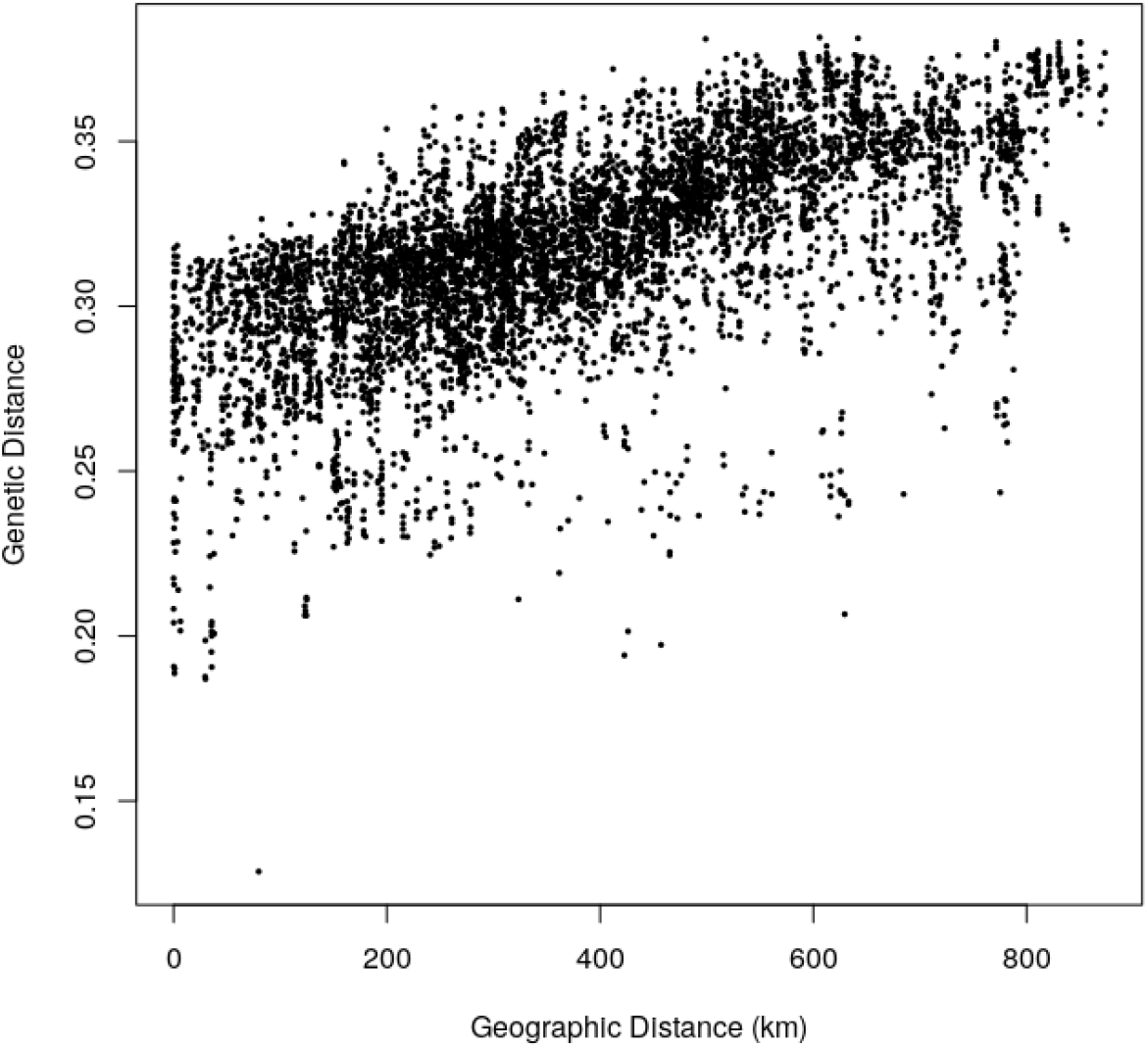
Isolation by distance in the leafcutting ant *Atta texana*. Genetic distance was calculated using method 2 in the adegenet’s dist.genpop function. This figure is identical to Figure 3, except the figure here is simplified and does not overlay the colored bars indicating the impact of ecological differences (temperature, precipitation, fungus-type) shown in Figure 3.

**FIGURE S5.**
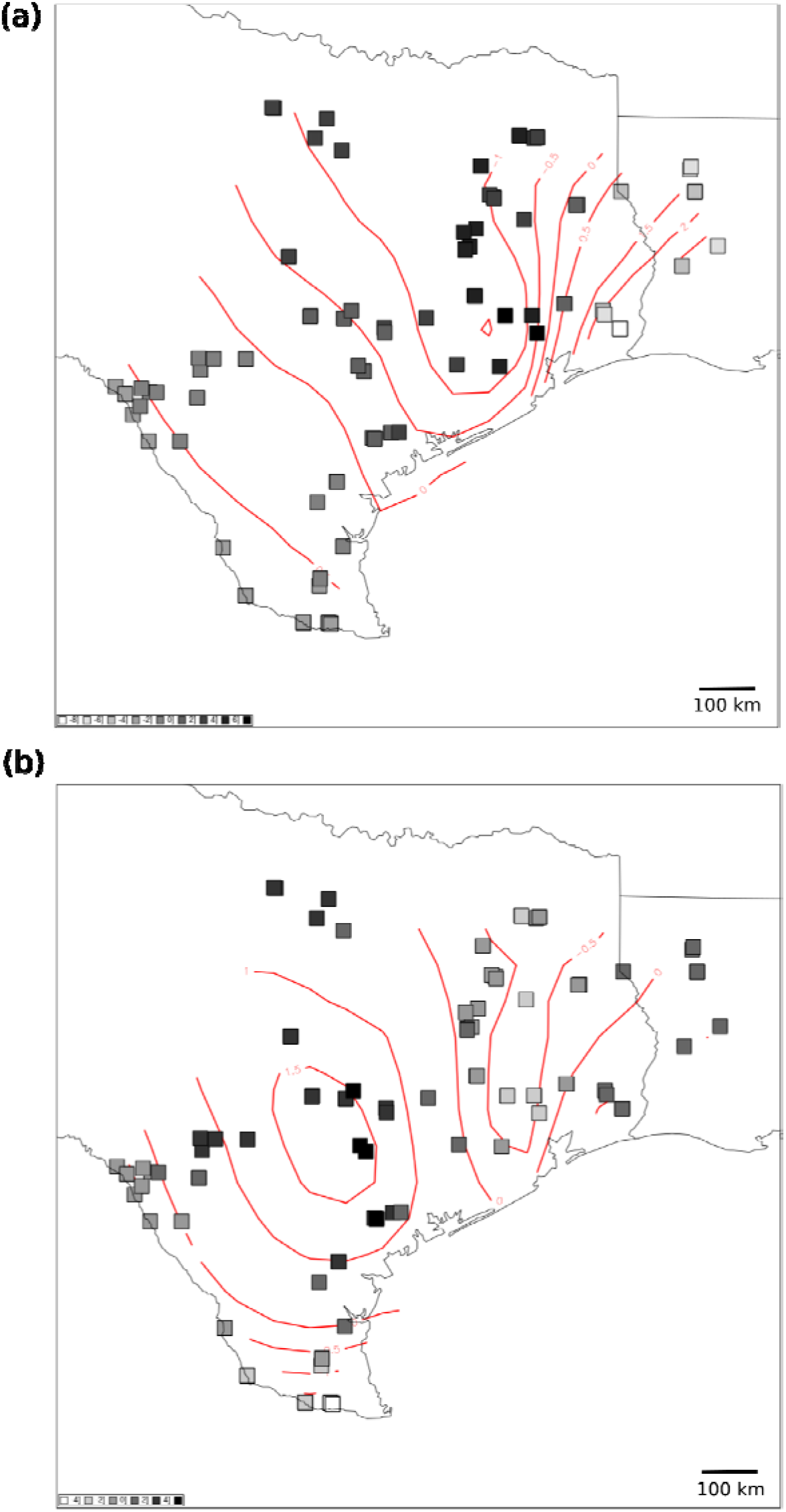
Global scores from the (a) second and (b) third axis of variation in the sPCA analysis of the leafcutting ant *Atta texana.* Genetic similarity is indicated by nests with similar shading. Contours represent the spatial distribution of global sPCA scores.

**FIGURE S6.**
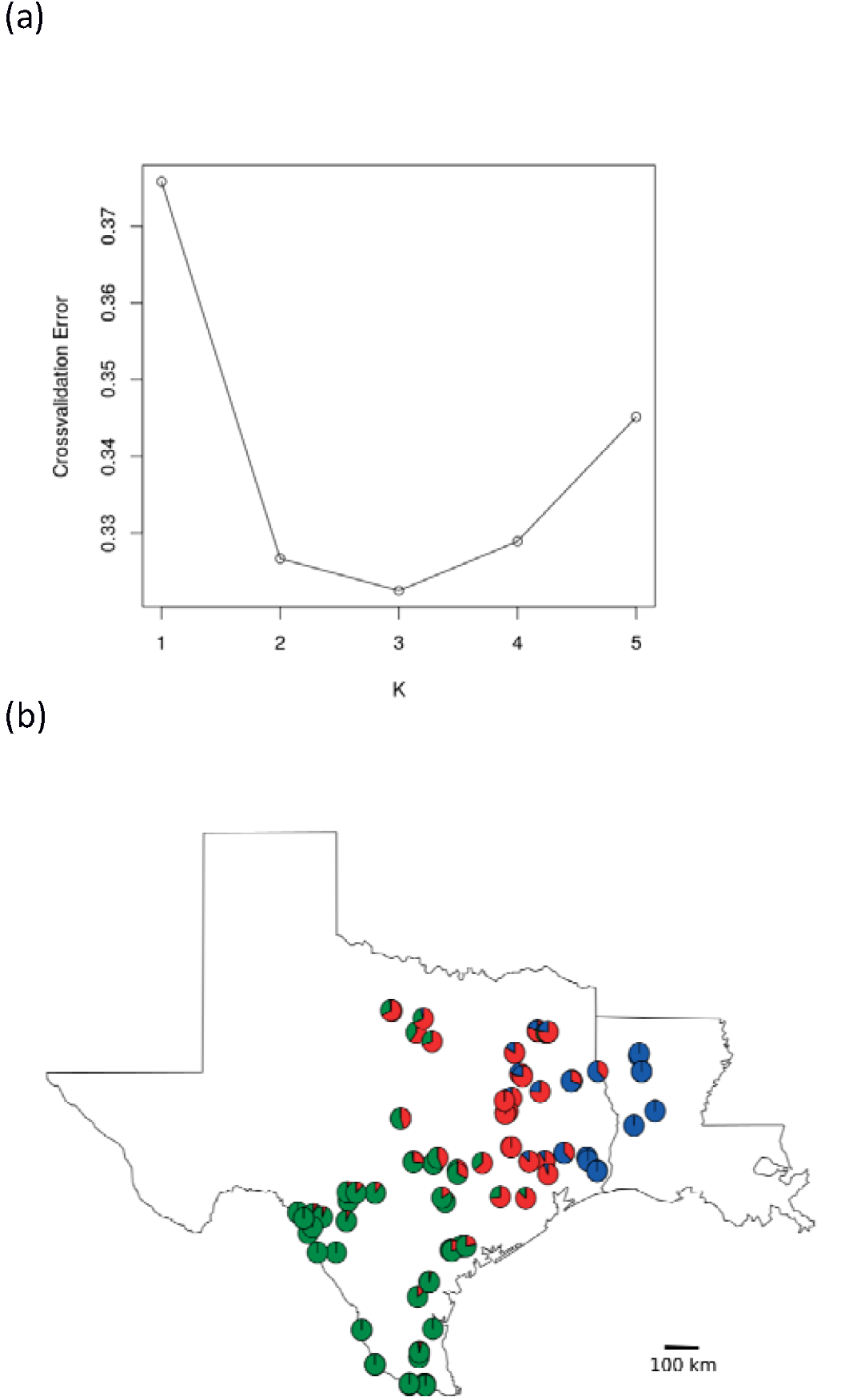
ADMIXTURE analysis of the ant *A. texana*. (a) Output of the cross-validation analysis indicated K=3 as the most likely number of genotype-clusters. (b) Spatial distribution of admixture among clusters across the states of Texas (left) and Louisiana (right) in the USA, showing the proportional assignment for each ant genotype to the three inferred clusters in the colors green (southwest Texas population), red (central & north Texas), and blue (east Texas and Louisiana). Collections cover the entire range of *A. texana* in the USA.

**FIGURE S7.**
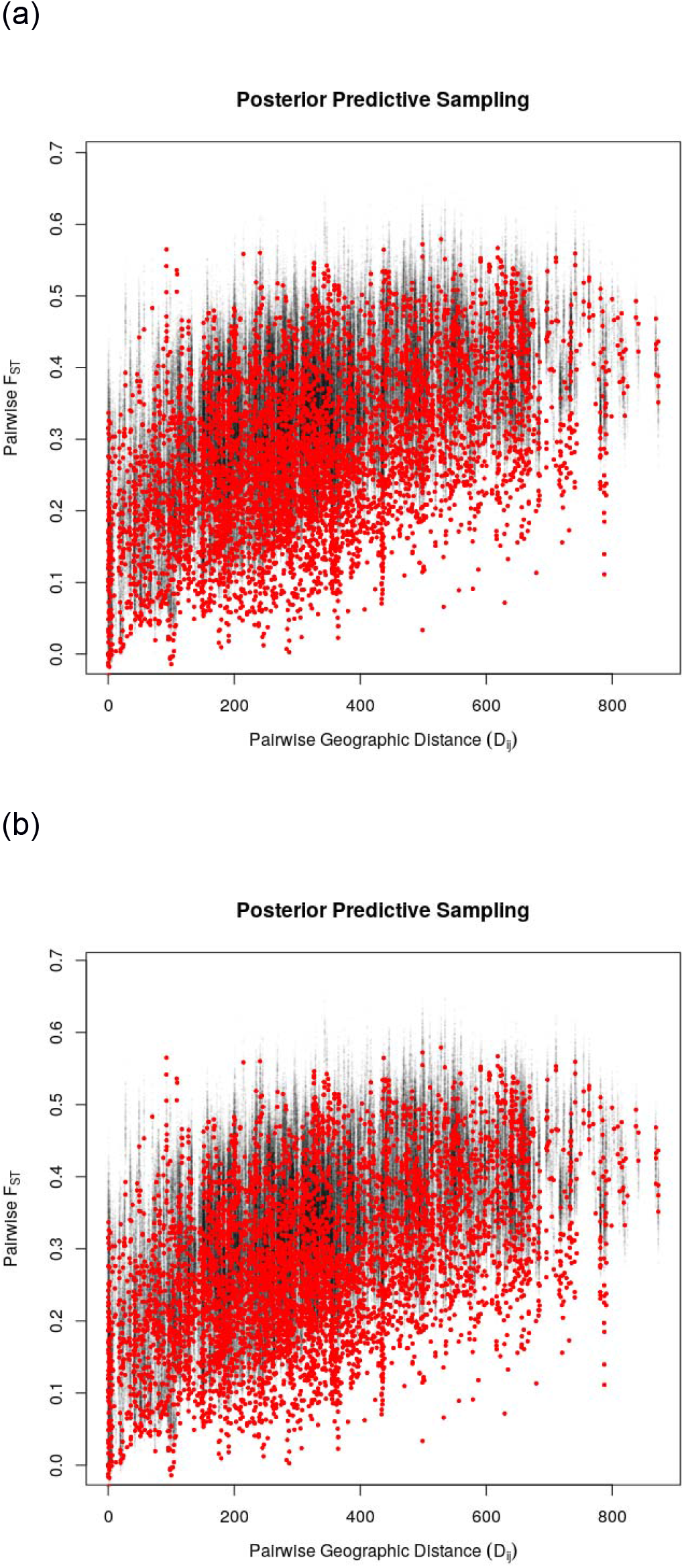
Posterior predictive sampling of BEDASSLE analysis with 100 simulated data sets for (a) the first replicate run and (b) the second replicate run.

